# The predominant binding mode of Palmatine to DNA

**DOI:** 10.1101/2024.09.17.613446

**Authors:** Marziogiuseppe Gentile, Francesco Talotta, Jean Christophe Tremblay, Leticia González, Antonio Monari

## Abstract

Palmatine is a protoberberine alkaloid, which may produce singlet oxygen under visible light irradiation and binds to DNA. The fact that singlet oxygen activation in palmatine may be triggered by environmental conditions, and in particular its interaction with nucleic acids, makes it a most suitable candidate for photodynamic therapy and DNA-targeted non-invasive anticancer strategies. Despite these remarkable properties the actual binding mode between palmatine and DNA has not been resolved, yet. Its elucidation has, indeed, led to contrasting hypothesis. In this contribution by using long-range molecular dynamic simulations and enhanced sampling approaches, we unequivocally identify that intercalation is the dominant binding mode of palmatine with DNA, from either a thermodynamic and kinetic point of view.

**TOC Graphic:** 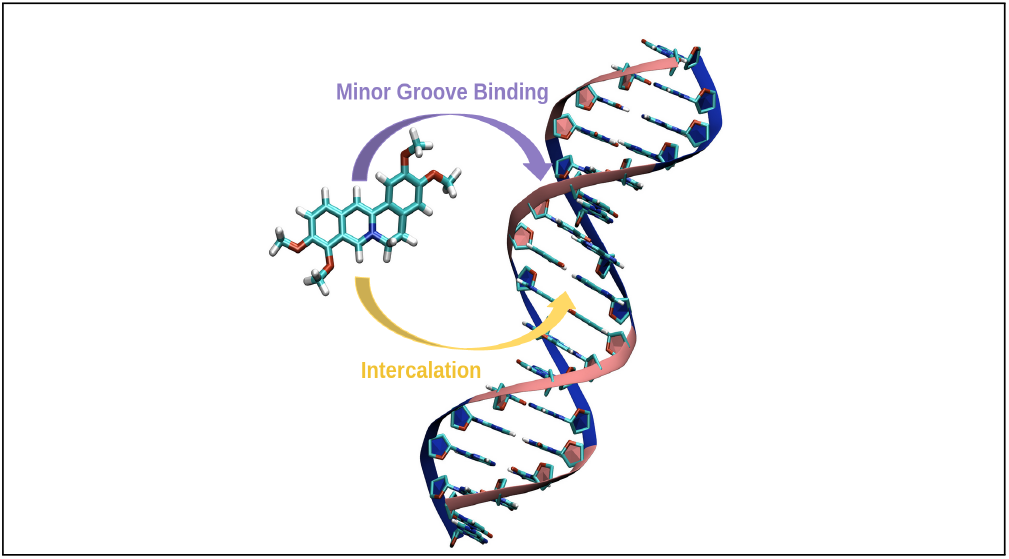

---

The exploration of small molecules that interact with DNA holds critical importance in drug development, especially in the field of targeted cancer therapies.^1–4^ Natural products, such as protoberberine alkaloids like palmatine (Figure 1a), have shown considerable promise due to their structural complexity and broad pharmacological effects.^5–7^ Understanding the specific ways in which these compounds bind to DNA, whether through groove binding or intercalation, is essential for harnessing their full therapeutic potential.

**Figure 1:**
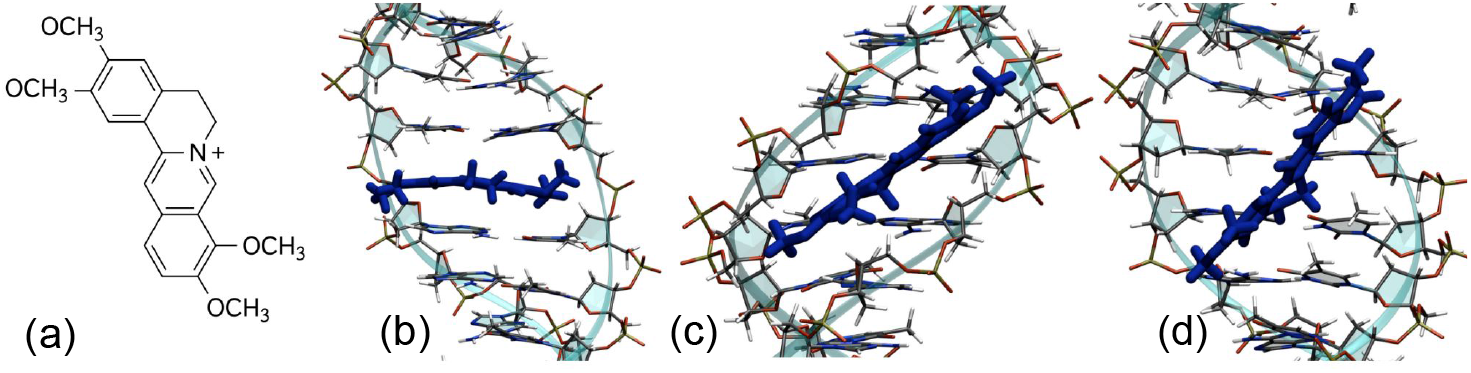
a) Structure of palmatine and representative snapshots of palmatine interacting with the poly(dA-dT).poly(dA-dT) DNA strand: intercalated (b), minor groove bound (c), or major groove (d). DNA atoms are represented in wire representation on top of ribbons. Carbon atoms are shown in grey, oxygen in red, hydrogen in white, and nitrogen in blue, palmatine is represented in licorice with blue color.

Palmatine, recognized for its inherent anticancer properties, is also suitable for photodynamic therapy (PDT) because it generates singlet oxygen ^1^O_2_ only when interacting with DNA.^8–10^ However, the interaction between palmatine and DNA, which is essential for modulating its photophysical behavior and therapeutic potential, remains a subject of debate.^11–15^ Indeed, since PDT relies on the activation of photosensitive drugs through light irradiation to produce reactive oxygen species and trigger cell death,^16–21^ its efficacy can be enhanced by compounds that exhibit specific and selective interaction modes with biological macromolecular structures, particularly DNA. Therefore, understanding the binding mode between palmatine and DNA is essential for advancing our fundamental knowledge of environmentally controlled photophysical processes and enhancing its potential for therapeutic applications.

Non-covalent interactions with DNA may lead to a different range of conformations, which include electrostatic binding, groove binding, and intercalation. Each of the binding modes present distinct characteristics and implications for molecular biology and drug design.^22–25^ Electrostatic binding occurs when cationic molecules, too large to fit within the DNA grooves or in the intercalation sites, bind to the solvent exposed surface of the double helix through Coulomb interactions with the negatively charged phosphate backbone.^26^ By contrast, groove binding is achieved by ligands which can accommodate inside the major or minor grooves encompassing the DNA double helix. Major groove binding involves larger molecules capable of forming van der Waals and hydrogen bond interactions. This binding mode may exploit the rather large groove size and is further favored by specific interactions, particularly with the thymine methyl groups, which are exposed within the groove.^27^ Smaller molecules, often characterized by a flat crescent shape that minimizes steric hindrance, preferentially lead to minor groove binding. This is partly due to the groove’s narrower and slightly deeper shape. Minor groove binding is particularly frequent in AT-rich regions,^28^ due to the higher flexibility of the groove. Intercalative binding is, instead, highly distinct in its structural characteristics, involving the insertion of planar, usually aromatic, moieties between adjacent DNA base pairs. This binding mode necessitates the separation of the base pairs and the creation of a hydrophobic pocket that facilitates the insertion. The driving force behind intercalation mode is due to the strong *π*-*π* stacking between the intercalator and the nucleic bases. This main contribution may also be enhanced by additional van der Waals, hydrogen bond, and, eventually, charge transfer interactions. Differently from the other binding modes, intercalation is also accompanied by strong local and global structural deformation of the DNA structure. These perturbations, including the elongation and partial unwinding of the helix, may impair critical biological functions, such as transcription, replication, and repair processes. Therefore, intercalative agents may also be exploited in synergistic therapeutic approaches to enhance the efficiency of potential chemotherapeutic agents, maximizing genetic instability in cancer cells.

The structure of palmatine, shown in Figure 1a, is characterized by a tetracyclic ring system and a net positive charge due to the presence of a quaternary ammonium group. Thus, it suggests the capacity for both intercalation and groove-binding. Yet, the general scientific narrative has not converged yet on a consensus regarding its dominant binding mode to DNA. This ambiguity not only hinders the rational design and optimization of palmatine-based lead therapeutic agents but also limits our ability to fully harness its potential in PDT applications. To study the interaction of palmatine with DNA, coherently with previous experimental studies,^9,12^ we considered two *in silico* built double-stranded DNA hexadecamers assuming a canonical B-form: 5’-AAAATTTTAAAATTTT-3’ (AATT) and 5’-AAGCTTTGCAAAGCTT-3’ (AGTC). We have then characterized their interactions with palmatine through all-atom molecular dynamics (MD) simulations. MD simulations can be used to explore the conformational space of a ligand-receptor complex and may help to identify the atomic details of the ligand-receptor interactions, providing further insights into the binding process.^29,30^ The initial systems have been constructed by a targeted manual positioning of palmatine within the DNA, building a starting conformation in which the ligand is bound to either the major or minor groove or is intercalated, Figure 1. The ensuing equilibrium MD simulations have shown that minor groove and intercalative binding are highly persistent, contrasting with the transient interactions observed in the major groove. This is also confirmed by the time evolution of the root-mean-square deviation (RMSD), which consistently indicates lower fluctuation for intercalative and minor groove binding modes, suggesting a stable interaction maintained throughout the simulation span. These results underscore the potential specificity and efficacy of palmatine for targeted binding. Detailed graphical representations of the time evolution of the RMSD are provided in Figure S1 of the Electronic Supporting Information (ESI) together with extended computational details.

Important conformational changes were observed in the DNA structure depending on the mode of binding. Intercalative binding induced a pronounced elongation and unwinding of the DNA helix. This alteration is consistent with the classical description of intercalation. Upon the ligand insertion between base pairs, an increase in the helical twist and the lengthening of the DNA axis are usually observed, to accommodate the ligand. Conversely, minor groove binding results in only minor structural perturbations. This observation again aligns with the expectation that minor groove binders typically do not significantly distort the DNA helix, preserving its overall canonical arrangement. To quantitatively assess the impact of these structural changes, we analyzed the DNA helical structural parameters using Curves+.^31^ The average values of rise and twist for the intercalative, minor groove, and ligand-free DNA are summarized in Table 1. The intercalative binding mode shows an increased average rise (3.7 Å vs. 3.2 Å) and decreased average twist (34° vs. 36°) across the DNA structure, compared to the unbound DNA. In contrast, the minor groove bound structure shows average rise and twist values much closer to those of the ligand-free DNA, confirming minimal structural perturbations.

**Table 1:**
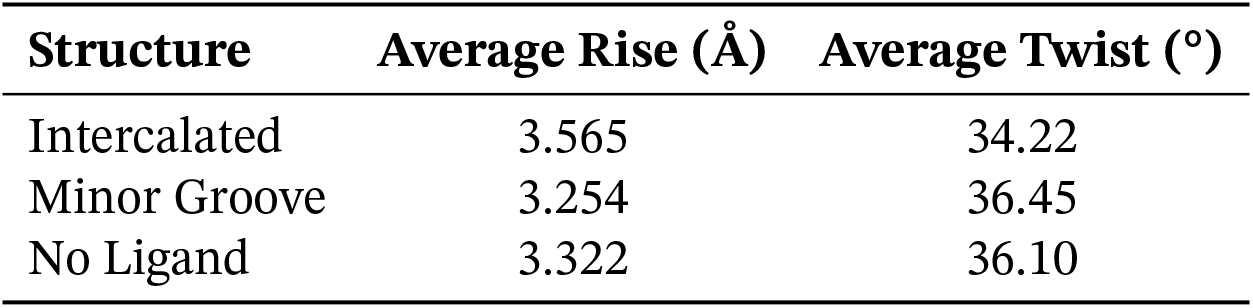
Average Helical Parameters for Different DNA Structures obtained from the MD simulations.

| **Structure** | **Average Rise (Å)** | **Average Twist (°)** |
| --- | --- | --- |
| Intercalated | 3.565 | 34.22 |
| Minor Groove | 3.254 | 36.45 |
| No Ligand | 3.322 | 36.10 |

In addition, we have also identified and analyzed the hydrogen bonding patterns and the dynamics of solvent interactions for the two persistent binding modes, i.e. intercalative and minor groove. Our MD simulations have revealed significant differences in the number and stability of hydrogen bonds between the DNA and the water molecules surrounding the ligand pockets for the two binding modes. This can be also appreciated by the analysis of the radial distribution function involving the solvent molecules which highlights the different interaction patterns. A consistent set of residues was used for these analyses, specifically considering the hydrogen bonds formed between the H4’, H5’, and H5” hydrogen atoms of the DNA sugar moiety and the oxygen atoms of the solvent water.

Upon intercalation, the number of hydrogen bonds per frame between DNA and the surrounding water reduces only slightly going from 2.43 to 2.29. This reduction is primarily due to the residues A9 and A25 (Figure 2a), which contribute to a loss of 0.13 hydrogen bonds. Conversely, for the minor groove binding mode, the number of hydrogen bonds per frame decreases more significantly to 1.60. this fact is mainly due to the displacement of water molecules from residues A9, A10, A26, and A27 (Figure 2b). Indeed, the analyzed hydrogen atoms on these residues develop strongers interaction with the *π*-system of the minor groove bound Palmatine. This might partially compensate the loss of hydrogen bonds between DNA and water, further stabilizing the binding mode.

**Figure 2:**
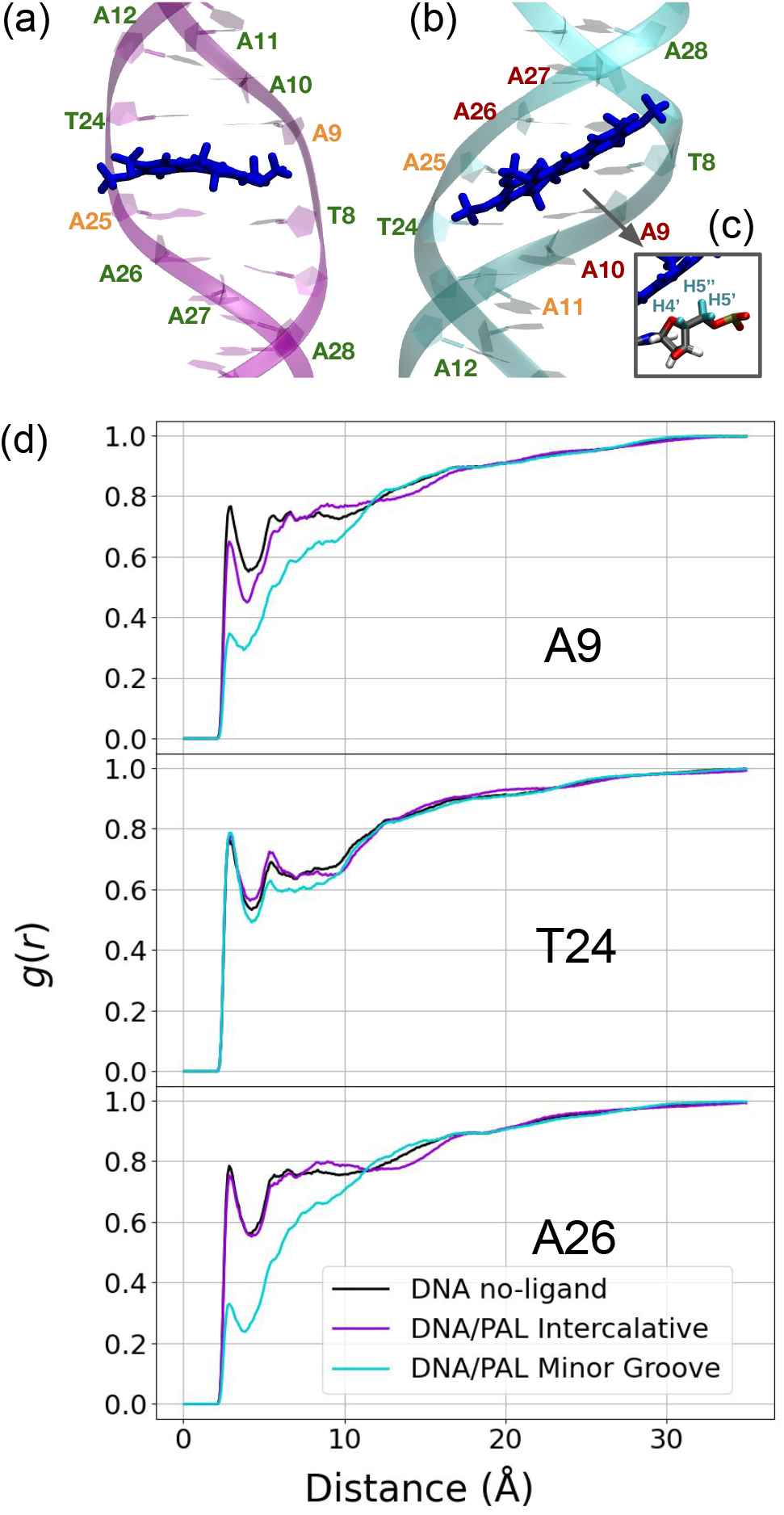
Representative snapshots for intercalative a) and minor groove b) binding modes of palmatine with DNA. c) Zoom of the H4’, H5’, H5” hydrogen atoms used or the calculation of RDF in the case of residues A9 d) RDF calculated by integrating the contributions of the H4’, H5’, H5” hydrogen atoms from the DNA strand with respect to the oxygen atoms of the solvent water molecules, for residues A9, A24, and A26. The corresponding DNA nucleotides are highlighted in panel a) and b) and are color-coded according to the strength of the hydrogen-*π* interaction (red: strong, orange: intermediate, green: weak).

These observations are supported by the Radial Distribution Function (RDF) analyses (Figure 2 and S2), which reveal a significant reduction of water density at short-distance for the minor groove binding mode, especially compared to intercalation.

These findings imply that the structural differences between intercalative and minor groove bindings influence the chemical environment of palmatine and DNA, in terms of hydrogen bonds and solvation. Specifically, the intercalative binding mode, despite significant DNA distortion, maintains a higher number of hydrogen bonds with the surrounding water compared to the minor groove binding mode, which disrupts the hydrogen bonding network more extensively by displacing water molecules. This effect is not entirely unexpected, as the DNA groove is exposed to solvent, requiring palmatine to displace water molecules in order to accommodate itself within the groove. On the contrary, intercalation mainly involves the hydrophobic, solvent inaccessible core of the double helix.

As summarized in Table 2, molecular mechanics Poisson–Boltzmann surface area (MM/PBSA) and molecular mechanics generalized Born surface area (MM/GBSA) methods have been used to estimate the enthalpic contributions to the binding free energy for the different DNA interaction modes. While at MM/GBSA level the energetic gain for the intercalative and minor groove binding mode is comparable, MM/PBSA previews a pronounced preference for intercalative binding. Indeed, an enthalpic contribution of -17 kcal/mol is estimated for the latter mode, significantly more favorable the one for minor groove and major groove bindings, amounting at -11 kcal/mol and -4 kcal/mol, respectivel.

**Table 2:** Binding enthalpies (kcal/mol) between Palmatine and DNA.

|  | Method / Mode | Intercalated | Minor Groove |
| --- | --- | --- | --- |
| Enthalpy | MM/GBSA | -33.21 | -34.57 |
|  | MM/PBSA | -17.29 | -11.47 |
| Free Energy | Umbrella Sampling | -10.30 | -5.40 |
|  | Isothermal Titration Cal.<sup>12</sup> | -7.03 | -5.36 |

The difference of the enthalpic contribution between intercalative and minor groove modes, approximately 6 kcal/mol, underscores the enhanced stability of palmatine when intercalated between DNA base pairs. The modest enthalpic gain observed for major groove binding aligns with the fact that this binding mode is not consistently maintained throughout the entire MD trajectory, indicating that it can be disrupted by thermal motion and entropic factors, which counterbalance the moderate energy stabilization. The enthalpic preference in the intercalative mode is likely due to more extensive molecular contacts, including stronger *π*-*π* stacking interactions and more favorable electrostatic stabilization, which are less pronounced in the minor groove. Likely the lore extended non-covalent interaction pattern is also counterbalancing for the energetic penalty due to the DNA deformation.

While MM/PBSA provides valuable insights on the thermodynamic ligand-DNA binding, the obtained binding energy is only semi-quantitative, and it does not offer direct information on the kinetic profile or the exact binding pathways. Indeed, although intercalative binding mode appears more stable from the MM/PBSA results, this binding mode could still be kinetically less favorable if a significant free energy barrier which could arise from the deformation of DNA, could be present. For this reason, we employed umbrella sampling, which allows exploring the free energy profile along a chosen collective variable. Most importantly, this method yields precise enthalpic and entropic contributions to the binding free energy through the calculation of the potential of mean force (PMF).

The umbrella sampling simulations have been conducted exploring, the distance from the centers of mass of palmatine and the four nucleobases that are closest to the binding site. For both binding modes, the collective variable *ξ* is represented in Figure 3.

**Figure 3:**
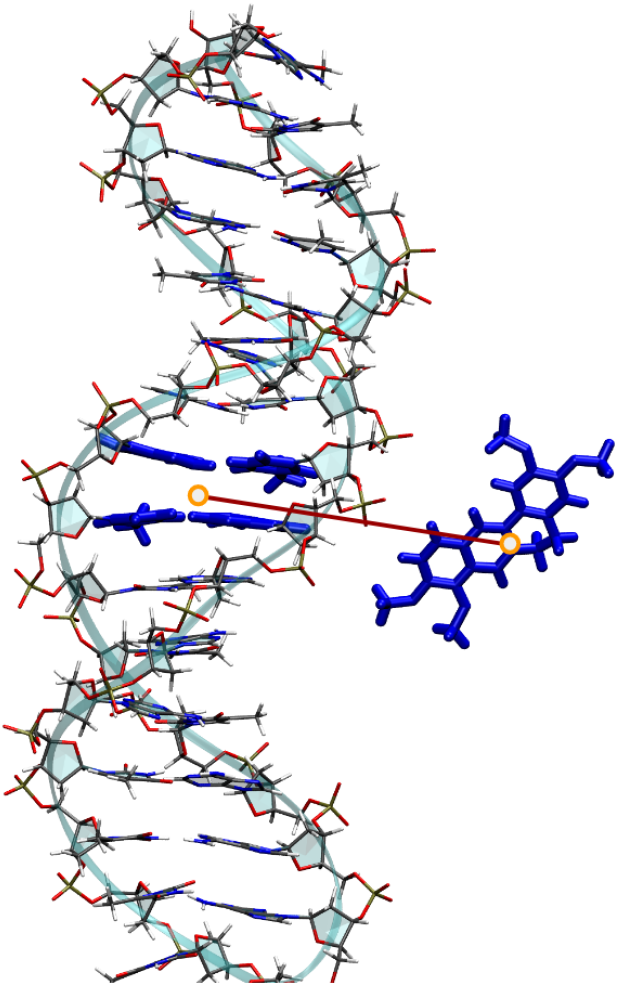
Graphical representation of the collective variable *ξ* used in the Umbrella Sampling procedure. The *ξ* variable involves the distance between the center of mass of Palmatine and the center of mass of the four nucleobases which are closest to the binding site

In the intercalated system, the nucleobases used to define the collective variable *ξ* are those directly involved in the formation of the intercalation pocket. For the minor groove-bound system, the nucleobases have been selected based on their closest proximity to the binding site.

As shown in Table 2, the calculated PMF yields a total binding free energy (ΔG) of -10.3 kcal/mol for intercalation and -5.4 kcal/mol for minor groove binding. These values compare well with experimental data obtained from isothermal titration calorimetry,^12^ which amount at -7.03 and -5.36 kcal/mol, for intercalation and minor groove binding, respectively. Interestingly, as also shown in Table 2, PMF and MM/PBSA preview a similar binding (fre) energy difference between the two modes amounting at about 6 kcal/mol in favor of intercalation. Importantly, as can be appreciated in Figure 5, no free energy barrier was observed in the PMF profiles for either binding modes. Similar results are also obtained for the AGTC DNA strand as reported in ESI (Figure S3 and Table S1). The suitability of the US profile was also checked assuring a good overlap between consecutive windows (Figure S4).

The absence of a free energy barriers for either intercalative and minor groove binding, can also be explained by the establishment of multiple, transient, non-covalent interactions between palmatine and the DNA backbone and phosphate groups, which may counterbalance the DNA deformation penalty. Indeed, in addition to the electrostatic interaction, palmatine is able to form hydrogen bonds and anion-*π* interactions involving simultaneously two adjacent phosphate groups, see Figure 4.

**Figure 4:**
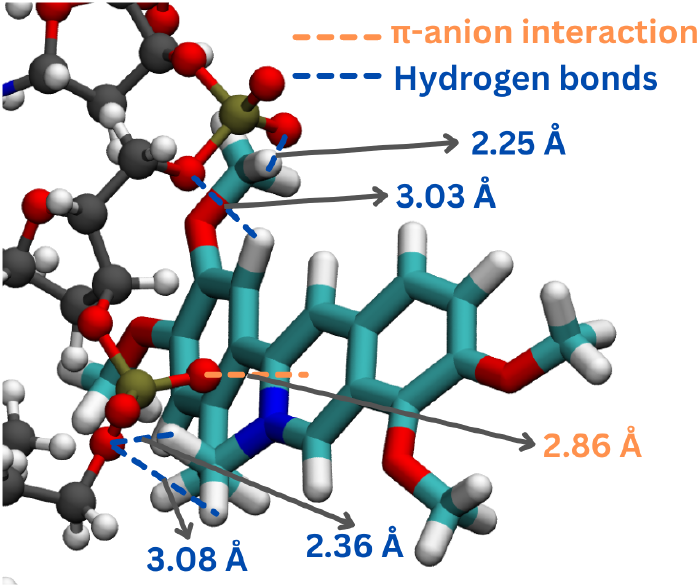
Representation of the non-covalent interactions, specifically hydrogen bonds and anion-*π* interaction, developing between palmatine and the DNA phosphate backbone. The average distance, in Å is also provided.

**Figure 5:**
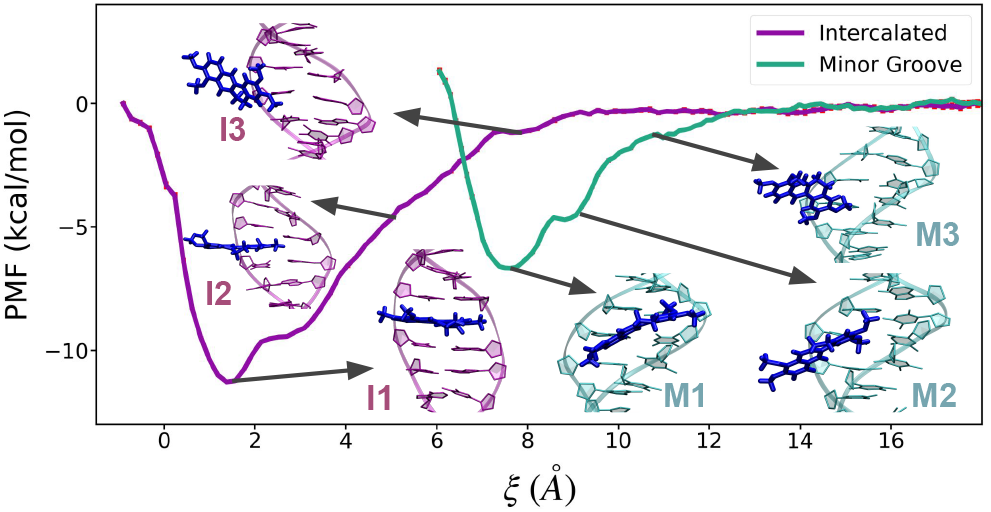
Potential of mean force (PMF) along the collective variable *ξ* describing the binding free energy of palmatine with DNA for the intercalative and minor groove binding modes. Selected configurations, representing different regions of the PMF are shown as inlays in purple (I1-I3) for intercalation and green (M1-M3) for minor groove, respectively.

From a mechanistic point of view, in the case of intercalation, as also illustrated in Figure 5, palmatine initially approaches the DNA from the major groove, and to subsequently come in close contact with the backbone phosphate (I3); it then progresses into a labile semi-intercalated state (I2) and eventually achieves the stable fully intercalated conformation (I1), aligning within the DNA structure. Similarly, for minor groove binding, palmatine initially moves towards the backbone phosphate (M3), to further sliding and rotating around the minor groove (M2), until it settles into the groove leading to the stable structure M1, and, thus, completing the binding process.

During the simulations, it was observed that palmatine experiences a stronger electrostatic attraction when approaching the DNA from the minor groove compared to the major groove. This enhanced attraction around the minor groove likely results from the closer proximity of the backbones of the two complementary DNA strands, allowing palmatine to simultaneously interact with both of them. Conversely, when approaching from the major groove, the backbone phosphates are at a greater distance, forcing palmatine to interact predominantly with only one DNA strand. This leads to a weaker attraction, which hinders its stable inclusion in the major groove, ultimately favoring intercalation. These observations suggest that the electrostatic interactions play a crucial role in guiding palmatine to its respective binding sites, with the spatial configuration of DNA significantly affecting both the accessibility and energetics of the binding site. However, the absence of kinetic barriers further indicates that once palmatine is near the DNA, it can readily adjust its position to achieve optimal binding, regardless intercalation or minor groove binding.

To validate the robustness of the previously outlined mechanisms, additional umbrella sampling simulations were performed to investigate palmatine behavior when starting from the major groove and pushing the ligand further into the groove. All these simulations consistently resulted in intercalation, confirming that the approach of palmatine to DNA from the major groove is a crucial factor in favoring intercalation.

Thus, our molecular dynamics simulations, MM-PBSA analyses, and free energy enhanced sampling calculation provide compelling evidence for a predominant intercalative interaction mode of palmatine into DNA. These methodologies underscore the pronounced enthalpic advantages of intercalative binding and highlight the stability and specificity of this interaction in the DNA helix structure. This detailed understanding may contribute to the development of molecular pharmacology of protoberberine alkaloids and offers important perspectives for the design and optimization of DNA-targeted therapies in cancer treatment. Our findings also confirm the potential of palmatine as an effective agent in PDT, due to its facile and barrierless intercalation,leveraging its ability to selectively generate reactive oxygen species when bound to DNA. In the future, we plan further computational studies to optimize singlet oxygen generation by palmatine near DNA. These studies will refine our understanding of spatial and electronic factors, including multichromophoric coupling, to enhance the efficacy and selectivity of palmatine-based therapies.

Further comparisons with closely related protoberberine alkaloids like berberine and coralyne could uncover a range of DNA binding affinities and mechanisms. Some derivatives might exhibit even higher affinities and greater therapeutic potential. Specifically, coralyne’s strong intercalative interaction with DNA makes it a promising candidate for enhancing anticancer effects beyond what palmatine can achieve. Establishing a robust protocol for detailing DNA interactions will facilitate a broader evaluation of these structural variations, ultimately guiding the optimization of therapeutic agents targeting DNA.

## Acknowledgement

The authors acknowledge the HPC resources of the University of Lorraine’s EXPLOR Mesocentre (https://explor.univ-lorraine.fr/). M.G. is grateful to Lorraine Université d’Excellence (LUE) for funding. A.M. thanks ANR and CGI (Commissariat à l’Investissement d’Avenir) for their financial support of this work through Labex SEAM (Science and Engineering for Advanced Materials and devices), ANR 11 LABX 086 and ANR 11 IDEX 05 02. Continuous support from the IdEx “Université Paris 2019” ANR-18-IDEX-0001, the Platform P3MB and the University of Vienna is gratefully acknowledged.

## Supporting Information Available

Computational Details, RMSD Analysis, RDF Analysis, US and MM/PB(GB)SA for AGCT DNA sequence, Distribution plots Umbrella Sampling

