## Supplementary Information for "The predominant binding mode of Palmatine to DNA"

### for

#### Contents

**S1 Computational Details** **S2**

**S2 RMSD Analysis** **S3**

**S3 RDF analysis** **S5**

**S4 US and MM/PB(GB)SA for the AGTC DNA sequence** **S5**

**S5 Distribution plots for Umbrella Sampling simulations** **S7**

**References** **S8**

### S1 Computational Details

Molecular dynamics (MD) trajectories have been run using the Amber 22 suite.[1,2] We considered two systems for DNA, i.e. the *in silico* built double-stranded 5'-AAAATTTTAAAATTTT-3' (AATT) and 5'-AAGCTTTGCAAAGCTT-3' (AGTC) hexadecamers in canonical B form. The DNA has been represented by OL21 force field.[3] The inital geometries were obtained with Amber NAB[4] code. Representative initial configurations of palmatine interacting with DNA for intercalation, minor groove and major groove binding have been manually constructed. The initial palmatine/DNA complexes have then been solvated in a OPC[5] (which couples well with the OL21 force field)[6] water box (buffer of 16.0 Å). The generalized Amber force field (GAFF2)[7] was used for modeling palmatine. Restricted electrostatic RESP[8] point charges have been obtained from quantum chemistry calculation using the established gaff protocol. The simulation box, which would be negatively charged due to the nucleic acid backbone, has been neutralized by adding the corresponding number of sodium cations (minus one due to Palmatine positive charge). After minimisation, thermalization and equilibration 3 independent trajectories of 100 ns each have been propagated in the isothermal and isobaric (NPT) ensemble at 300K and 1 atm. Pressure and temperature conservation have been enforced using the Langevin barostat and thermostat, respectively. The SHAKE algorithm[9] was used to constrain bond lengths involving hydrogen atoms, while the Particle Mesh Ewald (PME) approximation, with a cutoff of 10.00 Å, was employed to calculate long-range electrostatic interactions. Analysis of the trajectory focused on key structural and dynamical parameters, such as root mean square deviation (RMSD), radial distribution functions (RDF), and hydrogen bonding patterns, has been performed using CPPTRAJ.[10] The structures of the DNA and its deformation due to the presence of the ligand were analysed using Curves+.[11]

MM/PBSA and MM/GBSA[12–16] analyses have been performed on top of the equilibrium MD trajectories Amber to estimate binding energies. 13500 data points from the MD dynamics were used from each analysis, the first 1000 frames from the MD were discarded to avoid taking into account non equilibrated geometries.

To better assess the binding free energy for the stable poses we resorted to enhanced sampling approach using Umbrella Sampling (US). US simulations were performed using NAMD.[17] The distance from the centers of mass (COM) of palmatine to the COM of the four closest nucleobases in the binding site as collective variables. The collective variable was partitioned in windows of 1 Å, resulting in a total of 17 windows for the Intercalted system, and 11 for the Minor Groove binding. A force of 2.5 kcal/mol/Å$^2$ was used consistently. The initial geometries for each window were obtained from a Steered Dynamics in which palmatine is pulled away from the binding site to the bulk of the solution.

### S2 RMSD Analysis

The RMSD plots show that for either AATT and AGTC DNA sequences the Palmatine binds stably for the Intercalative and Minor Groove bindings, whereas the Major Groove binding result unstable. In the AATT Major Groove Binidng plot a flat RMSD is show from around 140 ns to 200 ns where the Palmatine migrate from the Major Groove and binds to the Minor Groove.

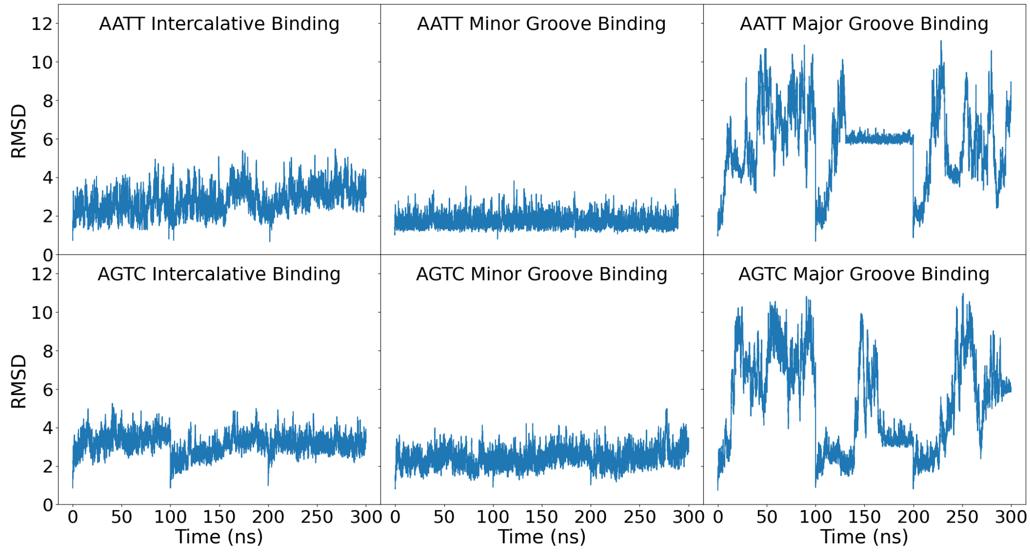


Figure S1: Root Mean Square Deviation (RMSD) of the palmatine geometry during Amber molecular dynamics for Intercalative, Minor Groove, and Major Groove Binding Mode, for both AATT and AGTC DNA sequences.

### S3 RDF analysis

In the plots are shown the RDF considering the DNA residues T8, A9, A10, A11, A12, T24, A25, A26, A27, A28 hydrogen atoms H4', H5' and H5"(averaged together) and Oxygen atoms of solvent water. The residues consider are displayed in the Figure 2a,b of the main text. Comparing the RDF plots it can be seen that when Palmatine binds to the Minor Groove some water molecules are displaced from it, whereas the RDF plot of the system with Palmatine Intercaled in DNA and DNA without any ligand show any substantial difference, highlighting instead that no water molecules is displaced.

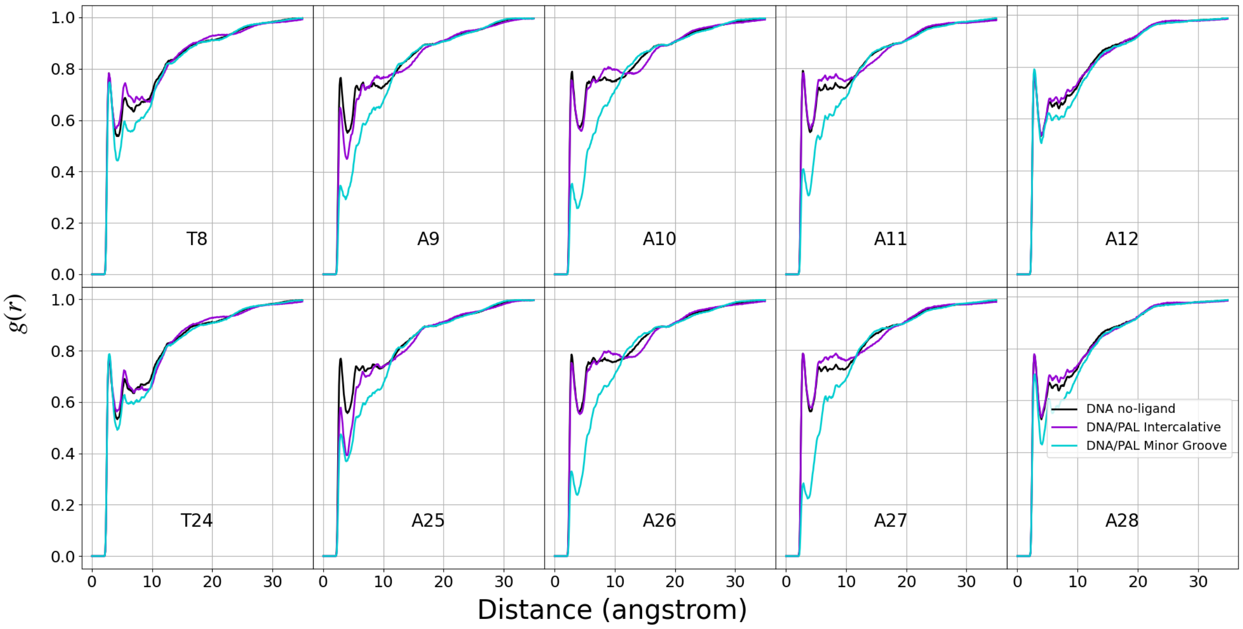


Figure S2: Radial Distribution Functions of the residues T8, A9, A10, A11, A12, T24, A25, A26, A27, A28, between DNA H4', H5' and H5" hydrogen atoms (averaged together) and the oxygen atoms of solvent water in presence or absence of palmatine.

### S4 US and MM/PB(GB)SA for the AGTC DNA sequence

Table S1 reports binding energies for Palmatine bound to the AGTC sequence, using either MM/PBSA, MM/GBSA, or Umbrella Sampling (US). Analogously to the AATT DNA sequence, Intercalative binding is found to be energetically favorable compared to Minor Groove binding for MM/PBSA and US. Further, US results indicate that the pathways for palmatine to the DNA strand for either Intercalative and Minor Groove binding are barrierless.

Table S1: Binding Enthalpies (kcal/mol) of Palmatine/AGTC DNA Complex

| Method / Mode | | Intercalated | Minor Groove |
|---|---|---|---|
| Enthalpy | MM/GBSA | -32.04 | -32.94 |
| | MM/PBSA | -14.53 | -10.72 |
| Free Energy | Umbrella Sampling | -16.77 | -6.91 |

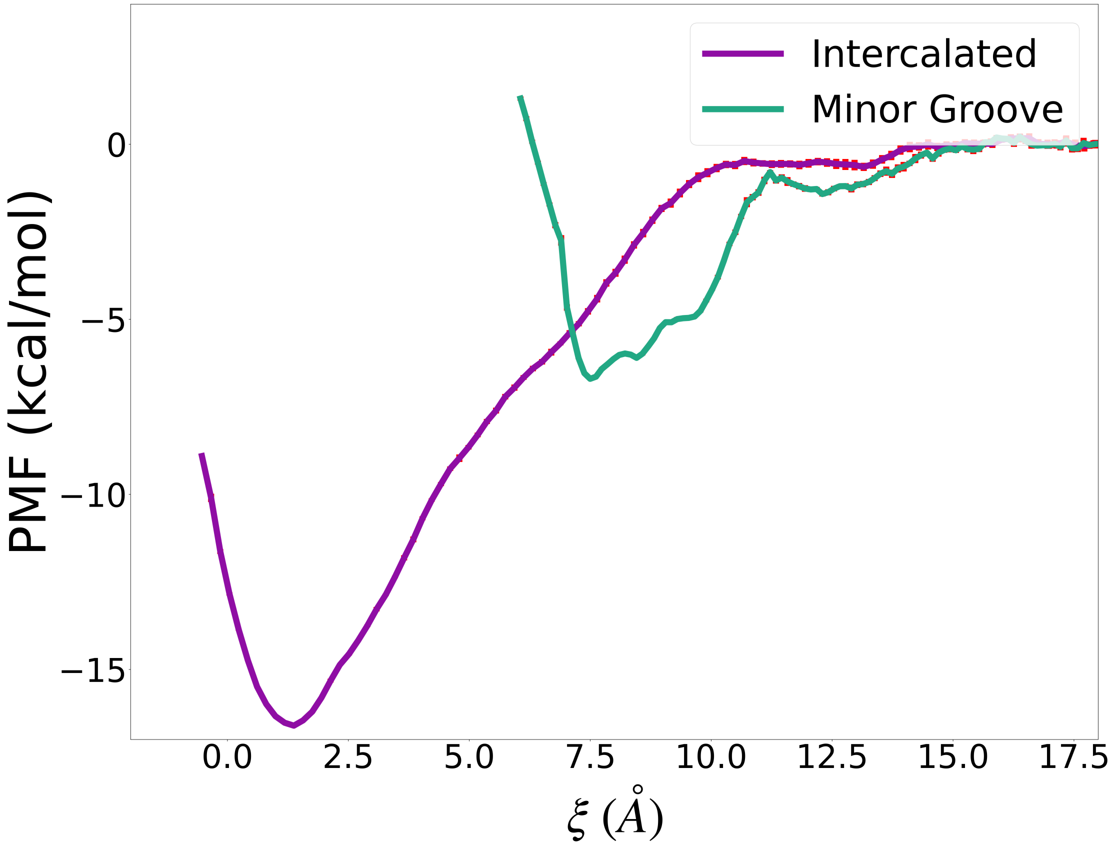


Figure S3: Potential of Mean Force for the Palmatine Intercalative and Minor Groove binding for the AGTC DNA sequence.

#### S5 Distribution plots for Umbrella Sampling simulations

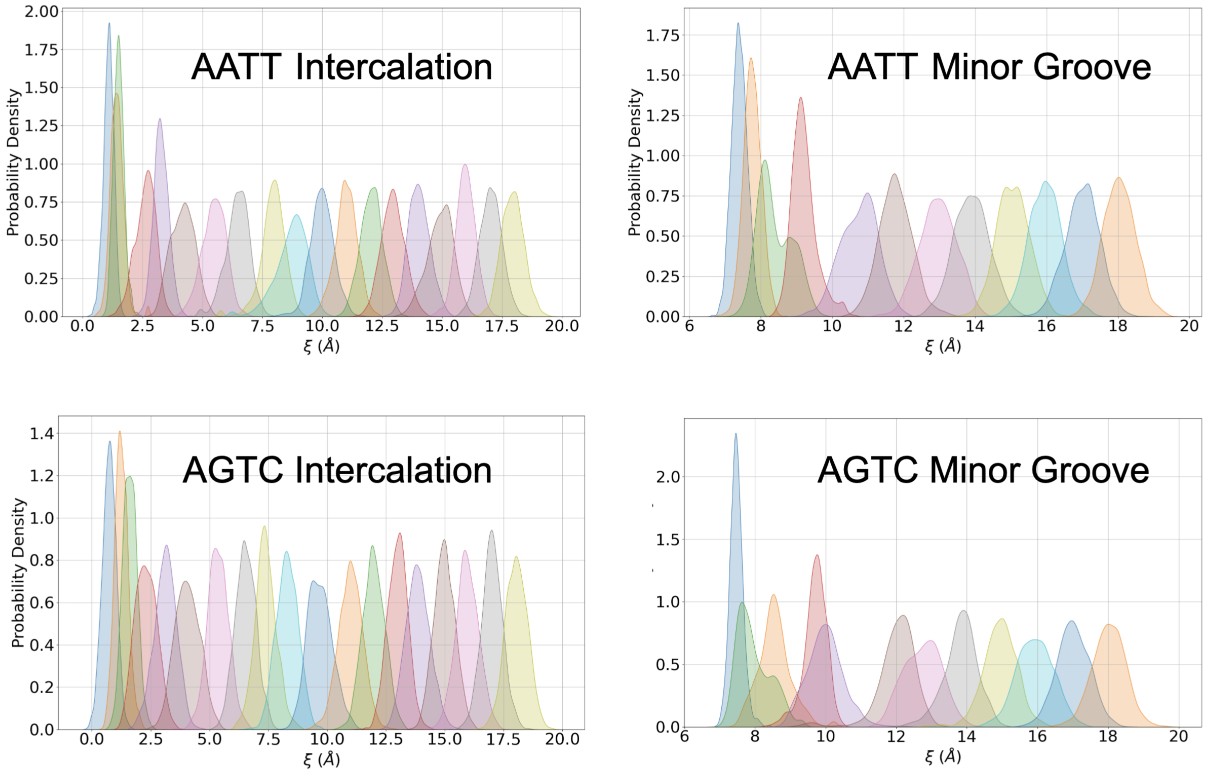


Figure S4: Geometry distribution plots for AATT and AGCT sequence for Intercalation and Minor Groove Binding showing an adequate windows overlap for the Umbrella Sampling procedure.
